# Renal carcinoma is associated with increased risk of coronavirus infections

**DOI:** 10.1101/2020.07.02.184663

**Authors:** Satyendra C Tripathi, Vishwajit Deshmukh, Chad J. Creighton, Ashlesh Patil

## Abstract

The current pandemic COVID-19 has affected most severely to the people with old age, or with comorbidities such as hypertension, diabetes mellitus, chronic kidney disease, COPD, and cancers. Cancer patients are twice more likely to contract the disease because of the malignancy or treatment-related immunosuppression; hence identification of the vulnerable population among these patients is essential. It is speculated that along with ACE2, other auxiliary proteins (DPP4, ANPEP, ENPEP, TMPRSS2) might facilitate the entry of coronaviruses in the host cells. We took a bioinformatics approach to analyze the gene and protein expression data of these coronavirus receptors in human normal and cancer tissues of multiple organs. Here, we demonstrated an extensive RNA and protein expression profiling analysis of these receptors across solid tumors and normal tissues. We found that among all, renal tumor and normal tissues exhibited increased levels of ACE2, DPP4, ANPEP, and ENPEP. Our results revealed that TMPRSS2 may not be the co-receptor for coronavirus in renal carcinoma patients. The receptors’ expression levels were variable in different tumor stage, molecular and immune subtypes of renal carcinoma. In clear cell renal cell carcinomas, coronavirus receptors were associated with high immune infiltration, markers of immunosuppression, and T cell exhaustion. Our study indicates that CoV receptors may play an important role in modulating the immune infiltrate and hence cellular immunity in renal carcinoma. As our current knowledge of pathogenic mechanisms will improve, it may help us in designing focused therapeutic approaches.

## Introduction

Coronavirus (CoV) disease-2019 (COVID-19) has been declared as a pandemic by the World Health Organization (WHO) after the outbreak of severe acute respiratory syndrome-CoV-2 (SARS-CoV-2) (Ng et al., 2020). The primary host for the SARS-CoV-2 has been identified as bats and the terminal host as humans (Khan et al., 2020). The related research revealed that SARS-CoV und SARS-CoV-2 share approximately 76% of amino acid identity (Wang et al., 2020). The primary symptoms include fever, dry cough, dyspnea, diarrhea, myalgia, headache, hyposmia and with common complications like acute respiratory distress (29%), acute cardiac injury (12%), and acute renal injury (7%) (Huang et al., 2020).

It has been well established that CoVs require the ACE2 or DPP4 receptors for entry into the host cells (Seys et al., 2018; Walls et al., 2020; Wrapp et al., 2020). The organs or the cells expressing ACE2 are more vulnerable to the CoV infection (Glowacka et al., 2011). According to Hoffmann et al., the viral entry in the host cell depends on the SARS-CoV receptor ACE2 for binding and requires the TMPRSS2 for priming and also relies on TMPRSS2 activity (Hoffmann et al., 2020). Therefore, it is speculated that other auxiliary proteins or co-receptors might facilitate the entry of CoVs in the host cells. These co-receptors or auxiliary proteins include TMPRSS2, ANPEP, ENPEP (Qi et al., 2020). Therefore, it has been proposed that organs representing the co-expression of co-receptors or auxiliary proteins such as TMPRSS2, ANPEP, or ENPEP for ACE2 and DPP4 are more susceptible for the viral entry replication and severity of the disease. The thought of extrapulmonary spread is not evitable due to the presence of these receptors and co-receptors. The prominent population with increased risk of virus infection are older patients and those associated with comorbidities such as hypertension, diabetes mellitus, chronic kidney disease, COPD, and cancers (Sun et al., 2020). Viral infection with comorbidities is responsible for higher mortality. According to Lee and colleagues, cancer patients are twice more likely to contract the infection than the normal population (Lee et al., 2020). Patients who received chemotherapy or surgery within the 30 days before the COVID-19 pandemic have more risk of infection than the patients who had not undergone chemotherapy or surgery (Sharma et al., 2020). According to an analysis of Italian patients published in March, 20% of those who died from COVID-19 in the country had active cancer (Liang et al., 2020). Notably, the guidelines for cancer patients during the COVID-19 pandemic focus on lung cancer patients undergoing active chemotherapy or radical radiotherapy, and on patients with blood cancers (Burki, 2020). The association of these receptors with the pathogenicity of the solid tumors is still to be solved.

In the present study, we investigated molecular profiling data of the various proteins required for the entry of the CoVs in normal tissues and cancer tissues. Immunological aspects of the study of pathogenesis cannot be overlooked. Therefore, we also explored an immune perspective concerning cancer. Understanding the usage of the multiplicity of receptors and co-receptors by the various CoVs can open new avenues for understanding the pathogenesis and development of intervention strategies.

## Results

### Expression profiles of CoV receptors in normal and tumor tissues

SARS-Cov-2 requires ACE2 as its receptor for host cell entry which is involved in various biological functions primarily in the renin-angiotensin system (Figure 1A). DPP4, ANPEP, ENPEP, and TMPRSS2 have also been proposed as co-receptors to initiate SAR-CoV-2 infection (Qi et al., 2020). STRING pathway analysis revealed a high confident interaction between these proteins, which have peptidase activity and are involved in angiotensin system, peptide metabolism, and viral entry into the host cell (Figure 1B and Table 1). Hence, we extracted the RNA and protein expression data of all these receptors in healthy tissues and solid tumors. We found enrichment of ACE2 at RNA levels in testis, small intestine, and kidney (Figure 1C). Along with ACE2, enrichment of RNAs for the other four co-receptors occurred in normal lung, mammary, liver, prostate, thyroid, head and neck tissues, small intestine, and kidney tissues (Figure 1C). The protein expression levels of ACE2 and co-receptors showed a similar expression pattern as to their RNA (Figure 1E). Renal tumors exhibited the highest expression of ACE2 receptor among solid tumor tissues followed by gastrointestinal cancers such as colorectal, pancreatic, and stomach cancer (Figure 1D). DPP4, ANPEP, and ENPEP RNA expression were also elevated in renal tumors compared to other cancer tissues. Of note, TMPRSS2 RNA expression was highest in prostate cancer tissues, whereas renal tumors featured among the lowest expressing tissues. TCGA data analysis also showed increased expression of all receptors except TMPRSS2 in renal tumor tissues as compared to their adjacent normal (Sup Fig 1). We found that protein level data was also in concordance to RNA expression data in solid tumors. Renal cancer expressed higher percent positivity of all receptors except TMPRSS2, which exhibited the highest percent positivity in prostate tumor tissues (Figure 1F,G).

**Figure 1:**
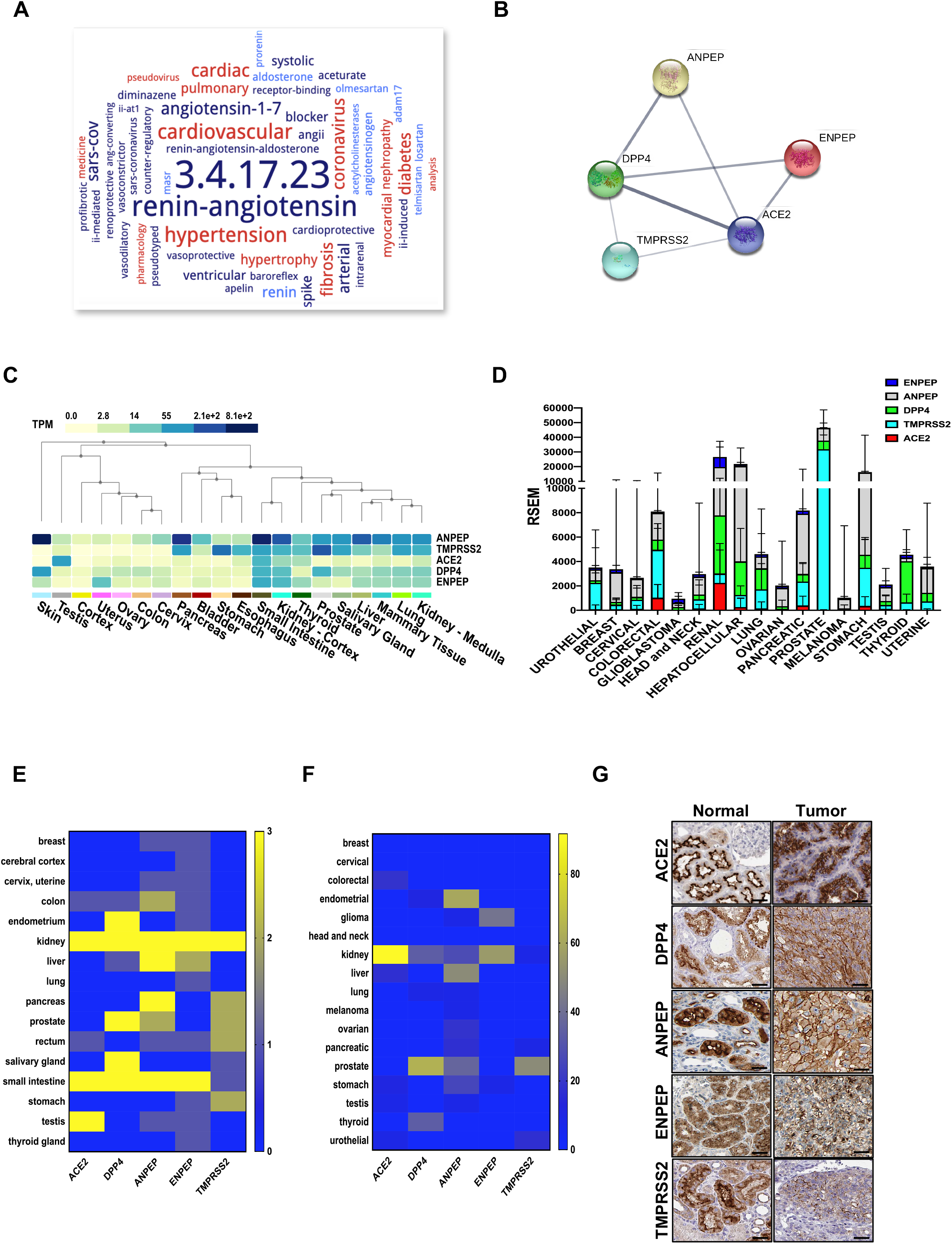
Expression of coronavirus receptors in human tissues. (A) The word cloud of various attributes associated with ACE2. (B) All coronavirus receptors were used with high confidence (0.8) evidence from experimental proteinLprotein interaction (blue lines) databases for STRING analysis. Each Node labeled with the encoding gene symbol represents a protein. (C) Heatmap of mRNA expression for coronavirus receptors in various human normal tissues and organs (GTEx dataset). (D) The mRNA expression profile of coronavirus receptors in different cancer tissues (TCGA dataset). (E, F) Heatmap for coronavirus receptors protein expression as analyzed by immunohistochemistry in (E) normal tissues and (F) cancer tissues (The Human Protein Atlas dataset). (G) Representative images of immunohistochemistry images of coronavirus receptors in normal and cancerous renal tissues (The Human Protein Atlas dataset). Scale bar = 50 um. ACE2, angiotensin-converting enzyme II. DPP4, Dipeptidyl-peptidase 4. ANPEP, Alanyl Aminopeptidase. ENPEP, Glutamyl Aminopeptidase. TMPRSS2, Transmembrane Serine Protease 2.

**Table 1:** Functional and molecular process related to coronavirus receptors

| GO Term | Functional process | FDR | Proteins |
| --- | --- | --- | --- |
| GO:0008238 | exopeptidase activity | 1.79E-07 | ACE2,ANPEP,DPP4,ENPEP |
| GO:0070011 | peptidase activity | 5.41E-07 | ACE2,ANPEP,DPP4,ENPEP,TMPRSS2 |
| GO:0004177 | aminopeptidase activity | 1.23E-06 | ANPEP,DPP4,ENPEP |
| GO:0008235 | metalloexopeptidase activity | 2.07E-06 | ACE2,ANPEP,ENPEP |
| GO:0001618 | virus receptor activity | 3.83E-06 | ACE2,ANPEP,DPP4 |
| GO:0070006 | metalloaminopeptidase activity | 5.08E-05 | ANPEP,ENPEP |
| GO:0004175 | endopeptidase activity | 0.00026 | ACE2,DPP4,TMPRSS2 |
| GO:0008270 | zinc ion binding | 0.0019 | ACE2,ANPEP,ENPEP |
| GO:0004252 | serine-type endopeptidase activity | 0.0023 | DPP4,TMPRSS2 |
| GO:0042277 | peptide binding | 0.004 | ANPEP,ENPEP |
| GO Term | Molecular process | FDR | Proteins |
| GO:0002003 | angiotensin maturation | 0.00025 | ACE2,ENPEP |
| GO:0006508 | proteolysis | 0.00025 | ACE2,ANPEP,DPP4,ENPEP,TMPRSS2 |
| GO:0008217 | regulation of blood pressure | 0.00025 | ACE2,ANPEP,ENPEP |
| GO:0016485 | protein processing | 0.00025 | ACE2,ENPEP,TMPRSS2 |
| GO:0046718 | viral entry into host cell | 0.00025 | ACE2,ANPEP,DPP4 |
| GO:0043171 | peptide catabolic process | 0.00047 | ANPEP,ENPEP |
| GO:0006518 | peptide metabolic process | 0.0019 | ACE2,ANPEP,ENPEP |
| GO:0010817 | regulation of hormone levels | 0.002 | ACE2,DPP4,ENPEP |
| GO:0001525 | angiogenesis | 0.0183 | ANPEP,ENPEP |

### Analysis of CoV receptors in Renal carcinoma

We observed that ACE2, ANPEP, ENPEP, and DPP4 gene expression was higher in renal papillary carcinoma (KIRP) and renal clear cell carcinoma (KIRC), whereas TMPRSS2 showed increased expression in renal chromophobe (KICH) (Figure 2a and Sup Fig 2a). We further analyzed the correlation of ACE2 with all four receptors in each renal cancer types. We observed a statistically significant correlation of ACE2 with DPP4 in KICH (p<0.01, ρ= 0.365), KIRC (p<0.001, ρ = 0.494), KIRP (p<0.001, ρ = 0.393) and also with ANPEP in KIRP (p<0.001, ρ = 0.277) (Figure 2B).

**Figure 2:**
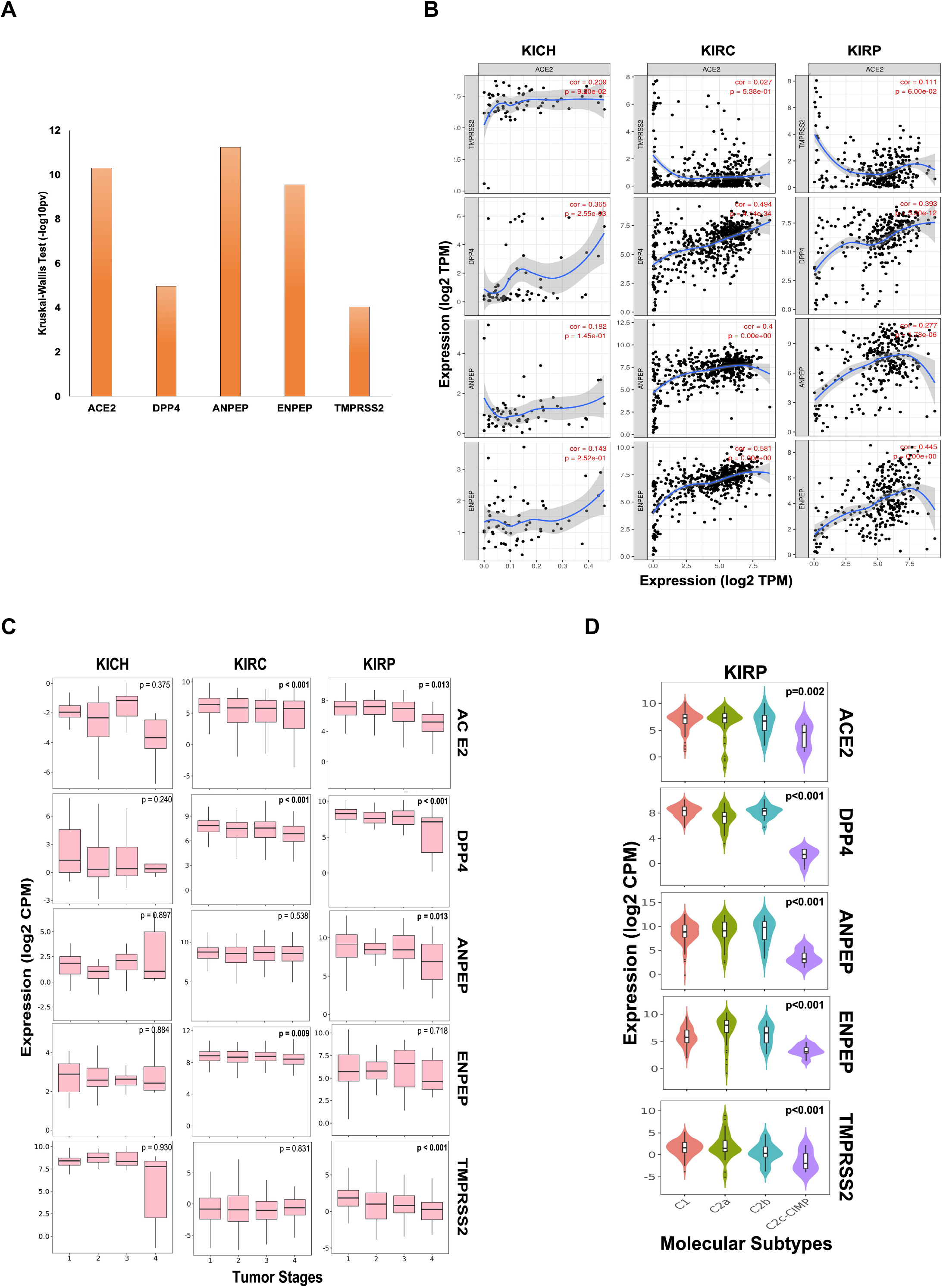
Coronavirus receptors in Renal carcinoma (A) The bar graph depicting a comparison of coronavirus receptor expression (TISIDB platform) (B) Correlation scatterplot of ACE2 with DPP4, ANPEP, ENPEP and TMPRSS2 (C) Boxplot for coronavirus receptors expression in different tumor stages (D) Violin plot for coronavirus receptors expression across various molecular subtypes in KIRP tumors. KIRP, Kidney renal papillary cell carcinoma.

Analysis of different tumor stages of renal carcinoma subtypes revealed a statistically significant (p<0.01) negative Spearman’s correlation of ACE2 and DPP4 with increasing stages of KIRC and KIRP tumors. However, TMPRSS2 exhibited very low expression levels, and also negatively correlated with tumor stages in KIRP tumor tissues along with ANPEP (ρ = −0.153, p<0.05). ENPEP only showed a significant (ρ = −0.112, p<0.01) negative correlation with tumor stages in KIRC tumors (Figure 2C). As molecular subtypes of KIRP are well defined, we also analyzed the expression pattern of these receptors in various molecular subtypes. We observed that all CoV receptors showed significantly (p<0.001, Kruskal–Wallis test) increased expression in C1, C2a, and C2b subtypes compared to C2c-CIMP subtype of KIRP (subtype with high DNA methylation) (Figure 2D).

### Association of CoV receptors with immune cell signature in renal carcinoma immune subtypes

As CoV infection leads to immunological consequences, we investigated CoV receptors' expression across the immune subtypes of renal carcinoma. The immune subtypes—C1 (wound healing), C2 (IFN-gamma dominant), C3 (inflammatory), C4 (lymphocyte depleted), C5 (immunologically quiet), and C6 (TGF-b dominant)—are each present across sizable subsets of KIRP and KIRC tumors (Ru et al., 2019). We observed that, except for TMPRSS2, the other four CoV receptors were highly significantly (p <0.001, Kruskal–Wallis test) expressed in C1-C4 and C6 immune subtypes of KIRC tumors. Whereas, immune subtype C5 had a higher expression of TMPRSS2 (Figure 3A and Supl Fig 2B). We also analyzed the correlation between the CoV receptor genes and immune cell signatures, for each of the 32 cancer types in TCGA. The CoV receptors tended to show a high correlation with immune signatures in most cancer types, though ACE2 and TMPRSS2 exhibited weaker correlations than other receptors (Supl Fig 2C). Further, we specifically analyzed ACE2, which is a primary SARS-CoV receptor and DPP4, highly correlated to ACE2 across the kidney cancer subtypes along with immune cell signatures (innate and adaptive immunity, inflammatory cytokines and chemokines). We observed that ACE2 and DPP4 exhibited increased expression in all molecular subtypes except KICH, CIMP subtype of KIRP, and CC-e.3 KIRC subtype (Figure 3B). We found that these receptors are highly correlated to the innate and adaptive immunity-related cells, as well as IL-10, IL-6, CXCL10, CCL2-CCL5, TGFB1 in KIRC tumors, whereas in KIPRP tumor tissues IL8, IL1A and TNF were highly correlated (Figure 3B). Our results revealed that, in KIRC tumors, the expressions of various markers of exhausted T cells (CD137, PD1, CTLA4) and immunosuppressive microenvironment (PDL1, PDL2) were also significantly (p <0.01, t-test) correlated to CoV receptors (Supl Fig 2D).

**Figure 3:**
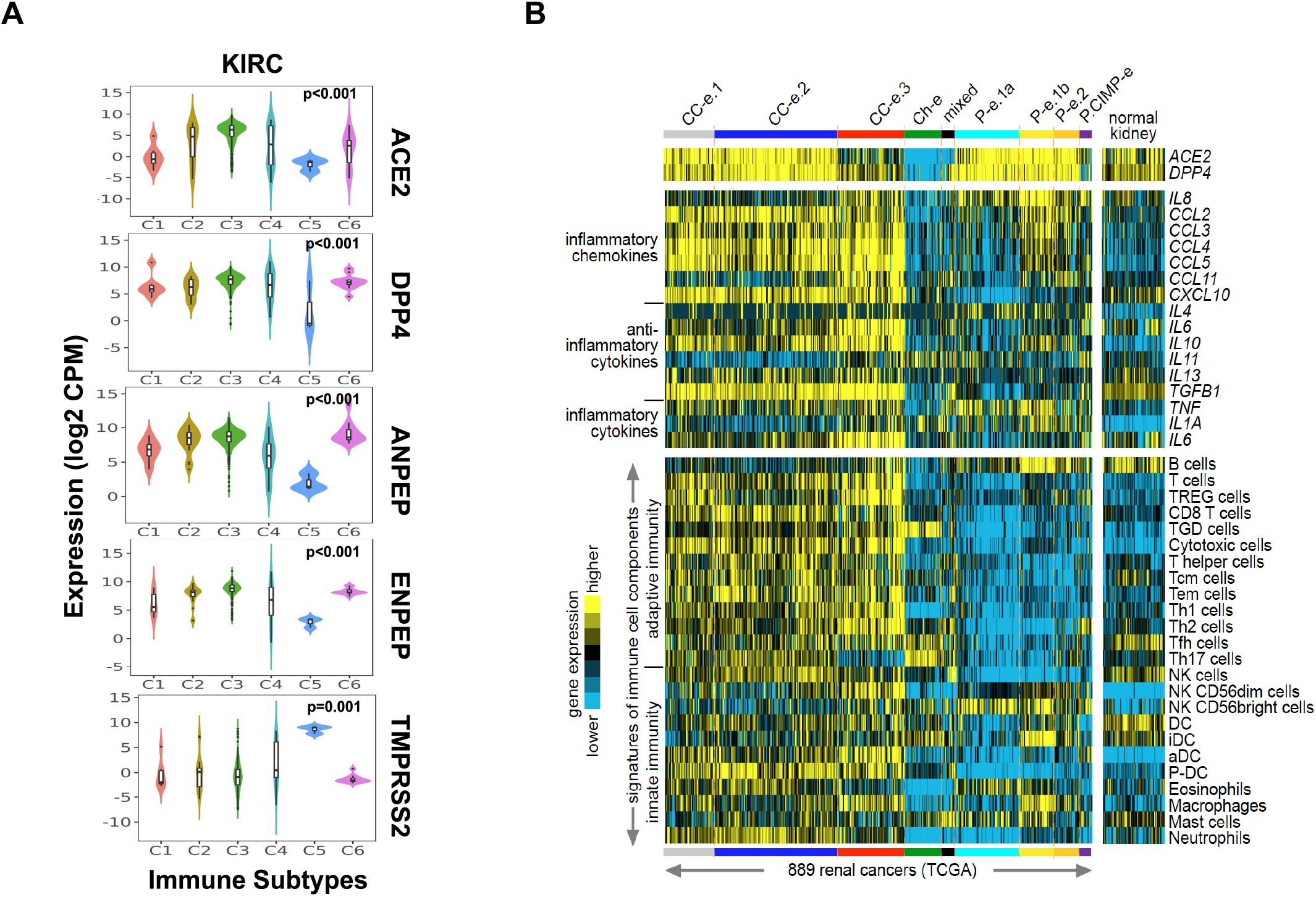
Immune signatures in renal carcinoma with coronavirus receptors expression (A) Violin plot for coronavirus receptors expression across various immune subtypes in KIRP and KIRC tumors. (B) Heatmaps of differential expression across renal cell carcinoma cases (from TCGA dataset (Chen et al., 2016)for genes encoding ACE2 and DPP4 (top), genes encoding selected chemokines and cytokines (middle), and gene expression-based signatures of immune cell infiltrates (bottom). Cases are ordered by molecular subtype as defined previously (Chen et al., 2016). Three of these subtypes—CC-e.1, CC-e.2, and CC-e.3—are enriched for clear cell RCC cases; four other subtypes—P-e.1a, P-e.1b, P-e.2, and P.CIMP-e—are enriched for papillary RCC cases; one subtype, Ch-e, is enriched for chromophobe RCC cases; and one subtype (“mixed”) is not enriched for any of the above. TREG cells, regulatory T cells; TGD cells, T gamma delta cells; Tcm cells, T central memory cells; Tem cells, T effector memory cells; Tfh cells, T follicular helper cells; NK cells, natural killer cells; DC, dendritic cells; iDC, immature DCs; aDC, activated DCs; P-DC, plasmacytoid DCs. KIRP, Kidney renal papillary cell carcinoma. KIRC, Kidney renal clear cell carcinoma.

### Immune infiltrate status and association with CoV receptors in Renal Cancer

We next investigated whether CoV receptor expression correlated with immune infiltration levels in renal carcinoma from TIMER. Notably, CoV receptors expression significantly correlated with infiltrating levels of various immune cells. In KIRC subtype, ACE2, DPP4, ANPEP, and ENPEP showed significant (p > 0.05), moderately positive Spearman’s correlation with infiltrating levels of B cells, CD8+ T cells, macrophages, neutrophils, and dendritic cells (DCs) (Figure 4 and Suppl Fig 3). ACE2 was only found to be significantly positively correlated to macrophage (r = 0.329, p = 1.04e-07) immune infiltrate in KIRP subtype (Figure 4A). On the other hand, DPP4 was significantly positively and negatively correlated to macrophage (r = 0.219, p = 5.13e-04) and B cell (r = −0.147, p = 1.85e-02), CD8+ T cell (r = 0.162, p = 9.17e-03) immune infiltrate in KIRP subtype (Figure 4B). We observed either negative or no correlation of TMPRSS2 expression with immune infiltrates in renal carcinoma subtypes (Suppl Fig 3A). We also found a significant correlation of ANPEP and ENPEP with immune filtrate in KIRC as compared to KIRP tumors (Suppl Fig 3B and 3C). Immune infiltrates of B cells, CD8+ T cells, Macrophages, and DCs significantly correlated with these receptors except TMPRSS2 in KICH tumors (Figure 4 and Suppl Fig 3). These results strongly suggest a variable host immune response to CoV infection depending upon the renal carcinoma subtypes.

**Figure 4:**
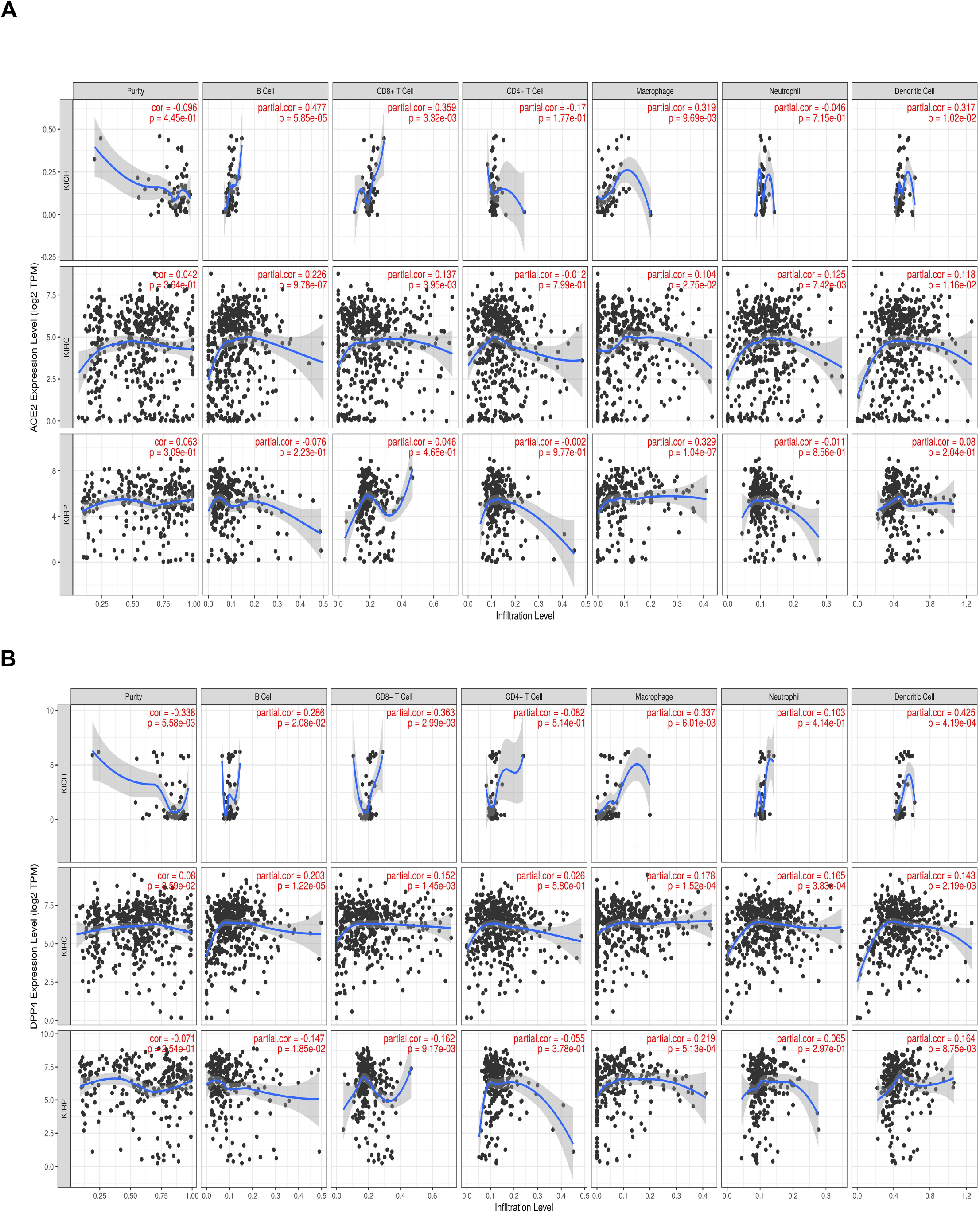
Correlation of coronavirus receptors and Tumor immune infiltrate (A, B) Correlation scatterplot of (A) ACE2 and (B) DPP4 with tumor purity and tumor immune infiltration of B cell, CD8+ T cell, CD4+ T cell, Macrophage, Neutrophil and Dendritic cell in renal carcinoma. KICH, Kidney renal chromophobe. KIRP, Kidney renal papillary cell carcinoma. KIRC, Kidney renal clear cell carcinoma.

## Discussion

There is an increased risk of coronavirus related fatalities in the subpopulation with any underlying health conditions or comorbidities. The risk factors for an increased susceptibility for CoV infection include, but not limited to, diabetes, heart disease, hypertension, chronic renal disease, chronic obstructive pulmonary disease, smoking and cancer (Guan et al., 2020). Cancer patients especially those undergoing chemotherapy and other anti-cancer treatment have increased risk of mortality due to CoV infection (Lee et al., 2020). The precise mechanism of increased severity in cancer remains unclear. In this study, we did landscape profiling of CoV receptors and co-receptors (viz ACE2, TMPRSS2, ANPEP, ENPEP, and DPP4) in various normal and cancer cells.

Our findings are in concordance with previous studies reporting ACE2 expression in stratified epithelial cells, colon, lung, liver, and kidney. Single-stranded RNA viruses can have multiple receptors for their host cell entry (Zhang et al., 2019). SARS-CoV utilizes ACE2, CD209, CLEC4G, and CLEC4M for its infection to host (Marzi et al., 2004; Yang et al., 2004; Gramberg et al., 2005; Wang et al., 2008). A recent study has suggested a few other receptors such as DPP4, ANPEP, ENPEP, and TMPRSS2 as co-receptors/auxiliary proteins to complement ACE2 in initiating SAR-CoV-2 infection (Qi et al., 2020). We analyzed the data available on the GTeX portal and observed the co-occurrence of these receptors in the small intestine and kidney at both RNA and protein levels.

Corona infection is a multiorgan diseased condition and not limited to the lungs. Few studies have reported low levels of ACE2, in lung parenchyma as compared to other normal tissues (Jia et al., 2005; Qi et al., 2020). Cell type specificity also exists for ACE2 expression, such as in lung expression is mainly in alveolar cells (Type 2 pneumocytes) and immune cells (B cells, T cells or myeloid cells) (Qi et al., 2020). In kidneys, most of the cells of proximal collecting tubules and proximal straight tubules exhibit increased expression. Cancer patients are always at higher risk of COVID-19 infection, and related case fatalities due to the compromised immune system increases comorbidities or treatment-related immunosuppression (Mehta et al., 2020; Tagliamento et al., 2020). We observed an increased expression of ACE2, DPP4, ANPEP, and ENPEP in renal tumor data available on TCGA followed by gastrointestinal cancers such as colorectal, pancreatic, and stomach cancer. This observation further raises the possibility that, along with lung cancer, patients with other types of cancer may also have elevated infection risk. Further analysis of renal carcinoma subtypes revealed that KIRP and KIRC tumors exhibit increased levels of DPP4, ANPEP, and ENPEP along with ACE2 receptors. We found that the expression of these receptors is inversely correlated to tumor stage and varies by molecular subtypes in renal carcinoma.

As host immune response is crucial to eradicating viral infection, immunological aspects related to these receptors cannot be overlooked. We observed that CoV receptors tend to show a high correlation with immune signatures in most cancer types. We further explored the possibility, whether CoV receptors are involved in modulating tumor immunity. Our analysis revealed for the first time that these receptors were correlated with immune cell infiltration in renal carcinoma. Our findings supported the immunoregulatory functions as follows: ACE2, DPP4, ANPEP, and ENPEP expression is closely related to the infiltration level of B cells, CD8+ T cell, macrophage, neutrophil, and dendritic cell. Cytokine storm has been well-defined and described with the pathogenesis of the disease. Chemokines are involved in several biological processes, such as the development of innate and acquired immunity, embryogenesis, and cancer metastasis (Coperchini et al., 2020). These chemokines along with cytokines recruit different immune cells (Poeta et al., 2019). We found that ACE2 and DPP4 were highly expressed and significantly correlated to the innate and adaptive immunity-related cells, as well as IL-10, IL-6, CXCL10, CCL2-CCL5, TGFB1 in KIRC tumors only. Upregulation of CXCL10 can enhance the levels of Tumor-infiltrating CD8+ T cell and natural killer cells (Humblin and Kamphorst, 2019; Kikuchi et al., 2019; Petty et al., 2019). T regulatory cells and macrophages get recruited by CCL5 (Wang et al., 2017; Walens et al., 2019). These results indicate that CoV receptors can play an important role in cellular immunity by modulating the immune infiltrate via cytokines and chemokines secretion.

Cancer cells also can evade the immune system by promoting T cell dysfunction and exhaustion (Jiang et al., 2015; Thommen and Schumacher, 2018). We analyzed the expression of inhibitory immune-checkpoint molecules in renal carcinoma subtypes to understand the dysregulation of the tumor microenvironment. Our results revealed that the expressions of various markers of exhausted T cells (CD137, PD1, CTLA4) and immunosuppressive microenvironment (PDL1, PDL2) are highly correlated to CoV receptors in KIRC tumors. Therefore, targeting these CoV receptors along with immune checkpoint inhibitors in coronavirus positive KIRC patients can be beneficial.

Our study has some limitations as our findings are based on correlation and associations drawn on analysis of data extracted from several public databases. Further experiments are warranted to confirm the role of CoV receptors in immune modulation of renal carcinoma.

In conclusion, our bioinformatics analysis revealed that renal carcinoma patients might be more susceptible to CoV infection. We found evidence that TMPRSS2 may not be the auxiliary protein for coronavirus infection in renal carcinoma. ACE2 and DPP4 increased expression in renal carcinoma tissues as compared to normal kidney. This association suggests that these patients are at increased risk of case related fatalities than healthy subjects. ACE2, DPP4, ANPEP, and ENPEP each associated with a high level of immune infiltration, inflammatory chemokines, cytokines and markers of an immunosuppressive microenvironment and T cell exhaustion in KIRC tumors. Our study indicates that CoV receptors may play an important role in modulating the immune infiltrate and hence cellular immunity in renal carcinoma.

## Methods

### Gene expression analysis

We downloaded RNA-Seq gene expression profiling datasets from the Genotype-Tissue Expression study (GTEx Consortium) for human normal tissues. The normalized RNA-Seq data in Transcripts per Million (TPM) was utilized for further analysis. We also downloaded RNA-Seq gene expression profiling datasets from The Cancer Genome Atlas (TCGA) (https://www.cancer.gov/aboutnci/organization/ccg/research/structural-genomics/tcga) (RSEM normalized) for human normal and cancer tissues.

### Protein expression analysis

The human protein Atlas database (https://www.proteinatlas.org/humanproteome/tissue) includes expression profiles of RNA and protein corresponding to ∼80% of the human protein-coding genes of specific tissues and organs. We downloaded the immunohistochemistry (IHC) data analysis of CoV receptors protein in cancer and normal tissues (PMID: 24309898). The score of IHC-based protein expression was determined as the fraction of positive cells defined in different tissues: 0 = 0–1%, 1 = 2–25%, 2 = 26–75%, 3 > 75% and intensity: 0 = negative, 1 = weak, 2 = moderate, and 3 = strong staining. We utilized the combined data of positive fraction and intensity, which is represented as high (3), moderate (2), low (1) or no (0) staining. The IHC data representation for normal tissues was presented as staining of the protein. For carcinoma tissues, we used the percentage positivity based on tumor tissues with high or moderate staining compared to low or no staining detected. Representative images of normal kidney and renal tumor tissues were also acquired from the same source.

For analyses of interactions among CoV receptors, the STRING database, which enables analysis for the structural and functional component of proteins (Szklarczyk et al., 2017)was used. The sources for establishing the interactions and enrichment of molecular, functional processes were Text mining, Experiments, Databases, Co-expression, Neighbourhood, and Co-occurrence, and 0.4 was set as the cut-off criterion.

### Immune infiltrate and subtype analysis

The Tumor Immune System Interaction database (TISIDB) (http://cis.hku.hk/TISIDB/) platform was used to analyze the correlation of ACE2 with other CoV receptors (DPP4, ANPEP, ENPEP, TMPRSS2) expression in different renal carcinoma subtypes. The correlation and association of Cov receptors with tumor stage, molecular subtypes, and immuno-subtypes of renal carcinoma were also analyzed using the TISIDB interface.

The Tumor Immune Estimation Resource (TIMER) (https://cistrome.shinyapps.io/timer/) platform comprised of the immune infiltrate data from the TCGA patients (Li et al., 2016, 2017) was used to investigate the association between CoV receptors expression and the infiltration level of B cell, CD4+ T cell, CD8+ T cell, neutrophil, macrophage and dendritic cell. “DiffExp” module was used to investigate the SPP1 expression between tumor and adjacent normal tissues across all TCGA tumors. The partial Spearman’s correlation and statistical significance after purity-correction were shown on the generated scatterplots.

### Immune signature enrichment in Renal carcinoma

To computationally infer the infiltration level of specific immune cell types using RNA-seq data from renal cell carcinoma samples from TCGA, as described previously (Chen et al., 2016), we used a set of 501 genes specifically overexpressed in one of 24 immune cell types (Bindea et al., 2013). We analyzed various immune signatures, including innate immunity, adaptive immunity, pro, and anti-inflammatory cytokines and inflammatory chemokines. The innate immunity marker genes included NK cells, Dendritic cells, Eosinophils, Macrophages, Mast cells, and Neutrophils. The adaptive immunity marker included genes such as B cells, T regulatory cells, T helper cells, and Cytotoxic T cells. We included cytokines like IL1A, IL4, IL6, IL8, IL10, IL11, IL13, TGFB1, and TNF. The chemokines were represented by CCL2, CCL3, CCL4, CCL5, CCL11, and CXCL10.

For pan-cancer correlation analyses, we used the set of 10224 RNA-seq profiles from 32 different cancer types as featured in our previous study (Chen et al., 2018), using the Bindea immune cell signature scores as computed for this study.

### Statistical Analysis

The correlation among CoV receptors expression and with immune infiltration level was determined by the TIMER interface using Spearman’s correlation analysis and statistical significance, and the strength of the correlation was determined using the following guide for the absolute value: 0.00–0.19 “weak,” 0.20–0.59 “moderate,” 0.60–1.0 “strong.” The association between the CoV receptors expression and molecular or immune-subtypes in renal carcinoma was analyzed by the TISIDB interface using the Kruskal-Wallis Test. For preparing the bar graph, the data was analyzed using GraphPad™ software (version 6.01, GraphPad Software, Inc., USA) and presented as mean ± SD. p-values < 0.05 were considered statistically significant.

