## Supplementary Figure Legends for "Renal carcinoma is associated with increased risk of coronavirus infections"

Dr Satyendra C. Tripathi

**Keywords: Coronavirus, Cancer, Kidney, ACE2, DPP4, TMPRSS2**

**Supplementary Figure 1**: Gene_DE module of TISIDB was used to analyze the differential expression between tumor and adjacent normal tissues. Distributions of coronavirus receptors expression levels across all TCGA tumors are displayed using box plots. Normal tissue data is displayed in gray columns when available. (*: p-value < 0.05; **: p-value <0.01; ***: p-value <0.001).

**Supplementary Figure 2**: (A) The bar graph for comparison of coronavirus receptor expression across renal cancer types (TISIDB platform) (B) Pan-cancer analysis, examining correlations between gene expression-based signatures of immune cell infiltrates (columns) and five different coronavirus related receptor genes (*ACE2*, *ANPEP*, *DPP4*, *ENPEP*, *TMPRSS2*), for each of 32 different cancer types represented in The Cancer Genome Atlas (TCGA). Each matrix entry represents the correlation between the given receptor gene expression and the given immune cell signature, for the given cancer type. Correlations by Pearson’s using log-transformed expression values. Purple, high correlation; cyan, low correlation. (C) Violin plot for coronavirus receptors expression across various Immune subtypes in KICH tumors. (D)

**Supplementary Figure 3**: Correlation of coronavirus receptors and Tumor immune infiltrate (A, B, C) Correlation scatterplot of (A) TMPRSS2, (B) ANPEP and (C) ENPEP with tumor purity and tumor immune infiltration of B cell, CD8+ T cell, CD4+ T cell, Macrophage, Neutrophil and Dendritic cell in renal carcinoma. KICH, Kidney renal chromophobe. KIRP, Kidney renal papillary cell carcinoma. KIRC, Kidney renal clear cell carcinoma.
