## Supplementary figures and images for "Renal carcinoma is associated with increased risk of coronavirus infections"

### Supplementary Figures

#### Suppl Figure 1

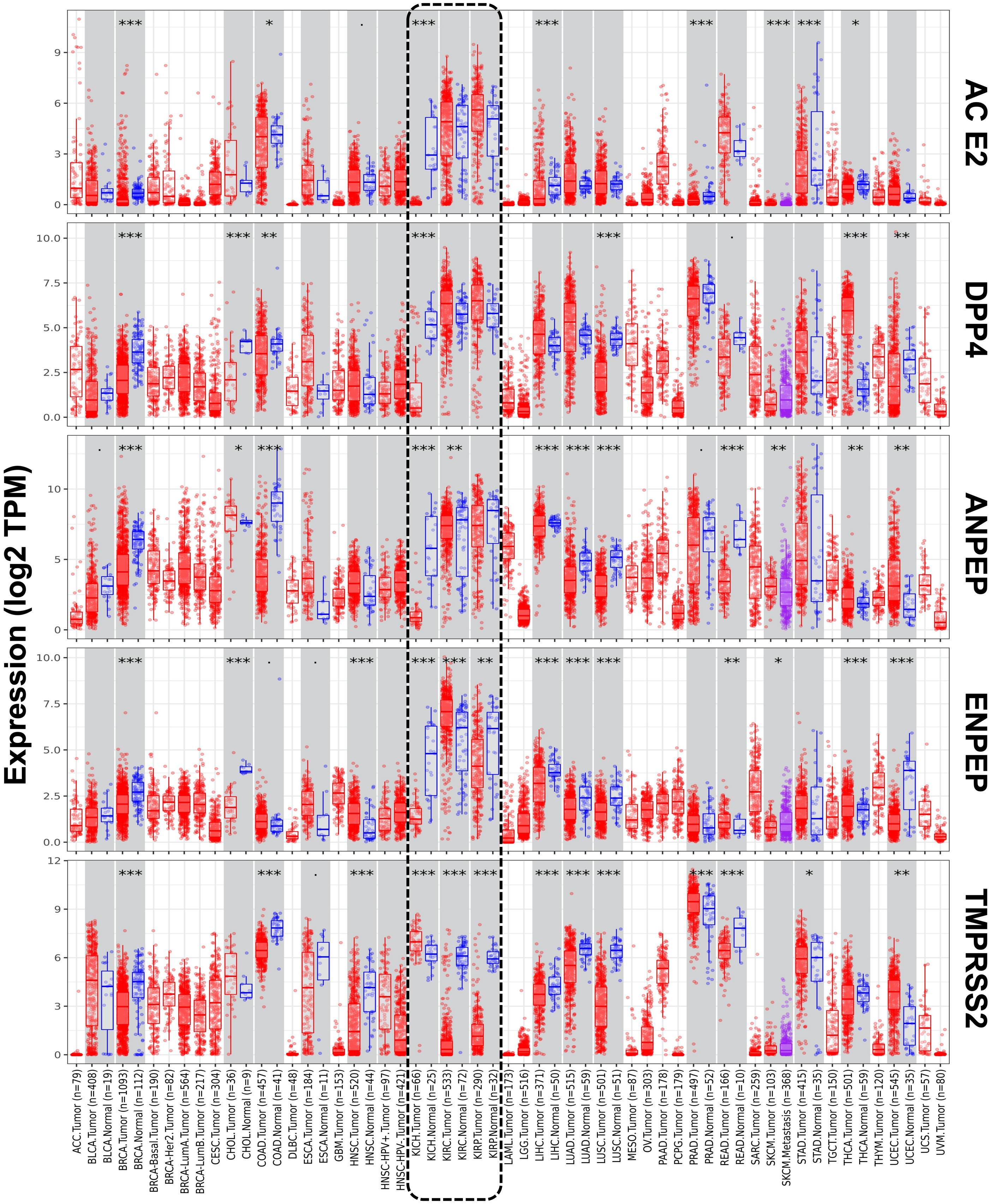

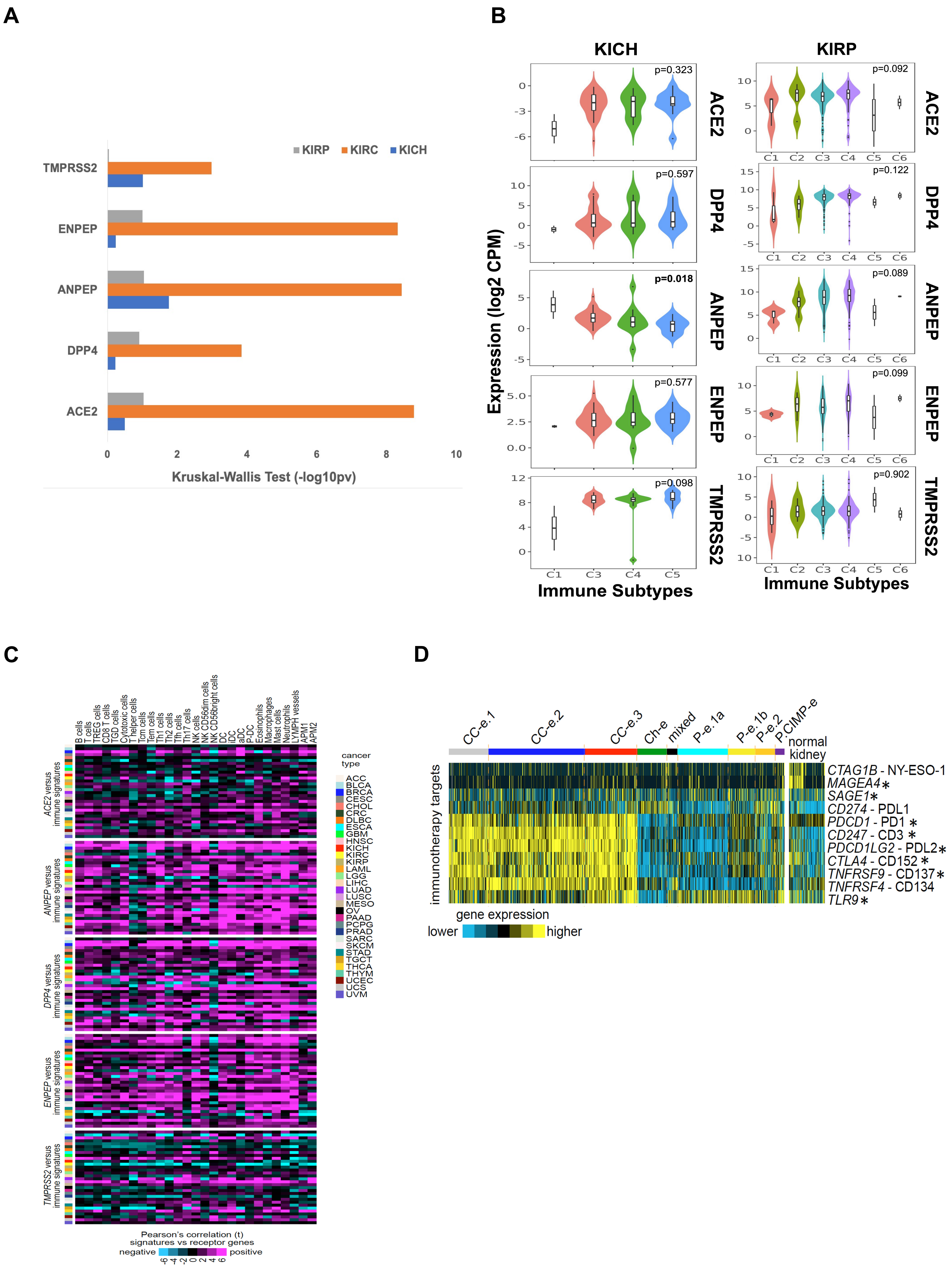

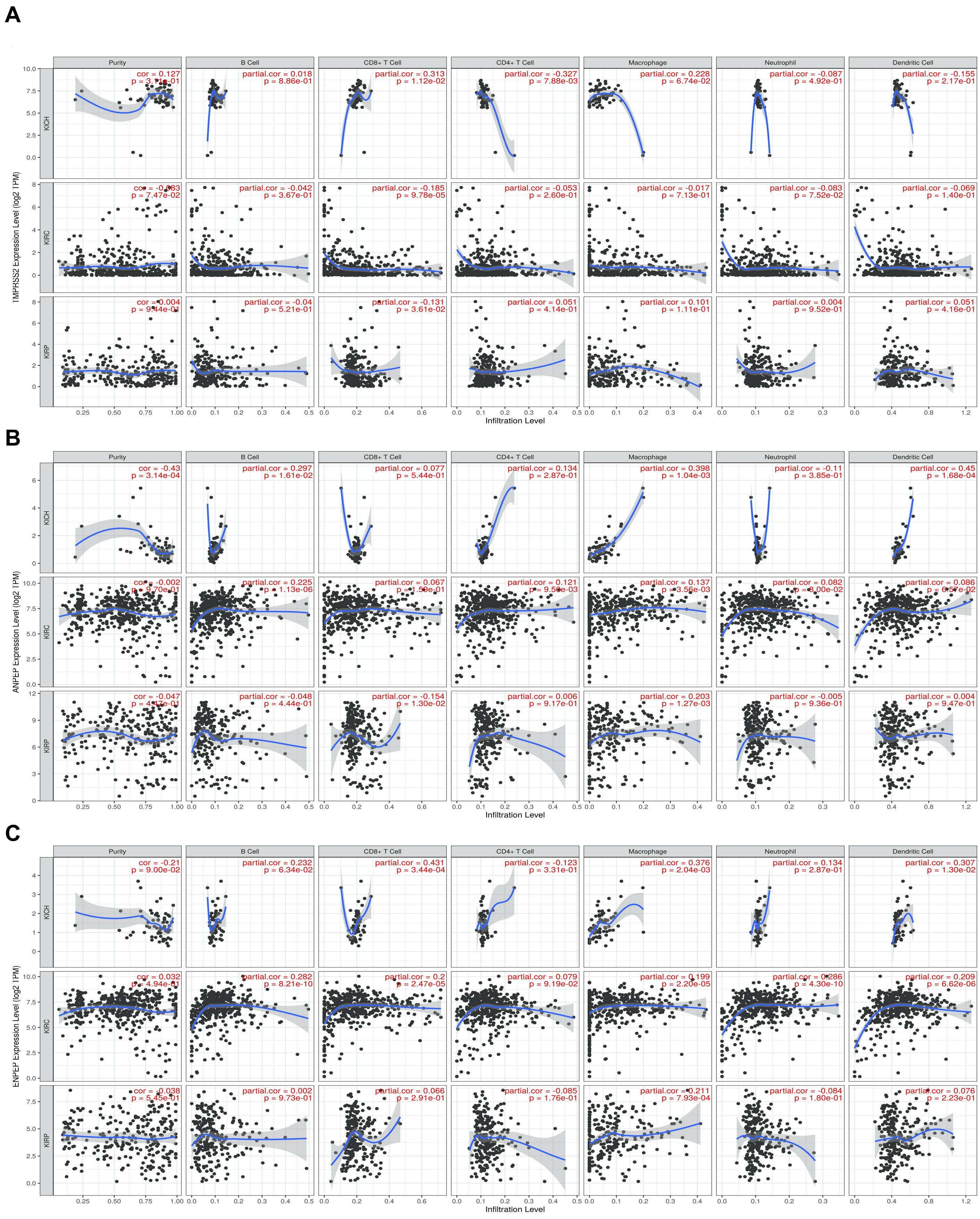
